# A DNA scaffold approach facilitates 5′ labelling of the SARS-CoV-2 RNA pseudoknot for smFRET investigation

**DOI:** 10.1101/2025.07.29.667457

**Authors:** Sarah P. Graham, Jemma K. Betts, Timothy D. Craggs, Mark C. Leake, Chris H. Hill, Steven D. Quinn

## Abstract

Single-molecule Förster resonance energy transfer (smFRET) studies of highly structured RNA molecules are often frustrated by issues with efficient dye conjugation. Here, we develop a DNA scaffold-based labelling strategy, and apply it to the frameshift-stimulating RNA pseudoknot from the SARS-CoV-2 genome. We prepare FRET-active reporters containing both Cy3 (donor) and Cy5 (acceptor) molecules and conduct measurements on freely diffusing single molecules, enabling the evaluation of conformational heterogeneity via smFRET population distributions. We identify that freely diffusing pseudoknots, modified at the base of stem 1, display a broad range of NaCl-dependent FRET states in solution, consistent with conformational freedom that extends beyond the static X-ray and cryo-EM structures. This work is a proof-of-principle demonstration of the feasibility of our DNA scaffold approach in enabling smFRET studies on this important class of biomolecule. Together, this work outlines new biochemical and biophysical approaches towards the study of RNA conformational dynamics in pseudoknots, riboswitches and other structured RNA elements.

## Introduction

Single molecule Förster resonance energy transfer (smFRET) spectroscopy has revolutionized the study of biological molecules, enabling conformational dynamics and structural heterogeneity of single molecules in solution to be revealed with nanometre scale resolution^1, 2^. In contrast to DNA and protein systems, for which smFRET has been well exploited and applied^3, 4^, applications to complex RNA systems have been generally under-developed. While structurally defined RNA systems such as hairpins^5^ have been routinely modified with fluorescent dyes and extensively studied in both ensemble and single molecule formats, single-molecule studies targeting RNA pseudoknots as structural motifs, though increasingly reported^6, 7, 8, 9, 10^, remain comparatively less explored. This is due in part to experimental challenges associated with the requirements to chemically conjugate fluorescent labels^11^ and the prohibitive costs of synthesising large RNA molecules with multiple internal dyes at specific locations. In this context, there are currently no FRET-based studies exploring the SARS-CoV-2 frameshifting element, an RNA pseudoknot.

The SARS-CoV-2 frameshift stimulatory element consists of three intercalated stems. Stem 1 (9 base pairs) is connected to stem 2 (5 base pairs) by a short loop 1 (3 nucleotides). Stem 2 stacks coaxially onto stem 1 via loop 2, while a third stem (stem 3) is derived from an extended loop 3 sequence. The result is a compact structure that resists ribosomal unwinding during translation of the viral genome^12, 13, 14, 15^. Single-molecule force spectroscopy of the pseudoknot under tension was recently reported, whereby single particles were tethered to duplex handles and attached to beads held in optical tweezers^16^. A variety of unfolding events were observed across a range of forces pointing towards conformational heterogeneity. Similarly, electron cryo-microscopy (cryo-EM) reconstructions^12, 13^, crystal structures^17, 18^, and molecular dynamics simulations^19^ indicate that the molecule can take on a variety of forms. Among these are unique topologies resembling knots, where the 5^′^ end passes through a junction of three helices, forming a ‘threaded’ structure commonly referred to as a “ring-knot”^12, 13, 19^. Interestingly, the observation of multiple conformations in the SARS-CoV-2 pseudoknot aligns with an emerging school of thought that –1 programmed ribosomal frameshifting is linked to structural heterogeneity within the stimulatory RNA element ^16, 20, 21, 22, 23, 24^. Solution-phase studies using small-angle X-ray scattering combined with molecular dynamics simulations^25^ also suggest notable deviations from the cryo-EM and X-ray structures—specifically more open and bent central ring structures were observed—highlighting the potential for conformational rearrangements under physiological conditions, and underscoring the need to further probe the structural freedom of the pseudoknot in solution. In this context, critical questions remain unanswered including (i) how many distinct conformational states does the pseudoknot adopt in solution? (ii) how dynamic are these states? and (iii) how do they respond to the local microenvironment?

Inspired by these insights, here we demonstrate the double labelling, single-molecule detection and smFRET characterisation of the RNA pseudoknot implicated in SARS-CoV-2 frameshifting. While previous work has mainly focussed on the structural characterisation of the pseudoknot in either a vitrified or crystalline state, we employed smFRET to probe the structural heterogeneity of single RNA molecules freely diffusing in solution. Importantly, smFRET allows direct access to the heterogeneity of single molecules and permits discrimination of sub-populations that may be otherwise hidden by the ensemble average, offering a unique perspective on the conformational landscape with nm-scale distance sensitivity to structural changes^26^. In contrast to X-ray crystallography, it is possible to conduct measurements in solutions more closely resembling physiological conditions: precipitants, high salt concentrations and extremes of pH can be avoided. By developing a labelling strategy to attach dyes to an occluded 5^′^ end, we were able to create double-labelled, FRET-active molecules that report on the distance between the 5_′_ end and the tip of stem 3. Our single molecule data reveal a broad range of NaCl-dependent FRET states, supporting the notion that the SARS-CoV-2 pseudoknot exists in a dynamic equilibrium of structures whose interconversion may play a role in frameshifting. We expect this approach to be broadly applicable to elucidating the conformational landscape of pseudoknots, riboswitches and other regulatory RNA elements, allowing for probing of structural responses to environmental factors, ligands and point mutations.

## Methods Materials

The SARS-CoV-2 RNA frameshifting element with internal fluorescent label (**Figure 1a**) was synthesised by Integrated DNA Technologies and purified by RNase-free high-performance liquid chromatography (HPLC).

**Figure 1.**
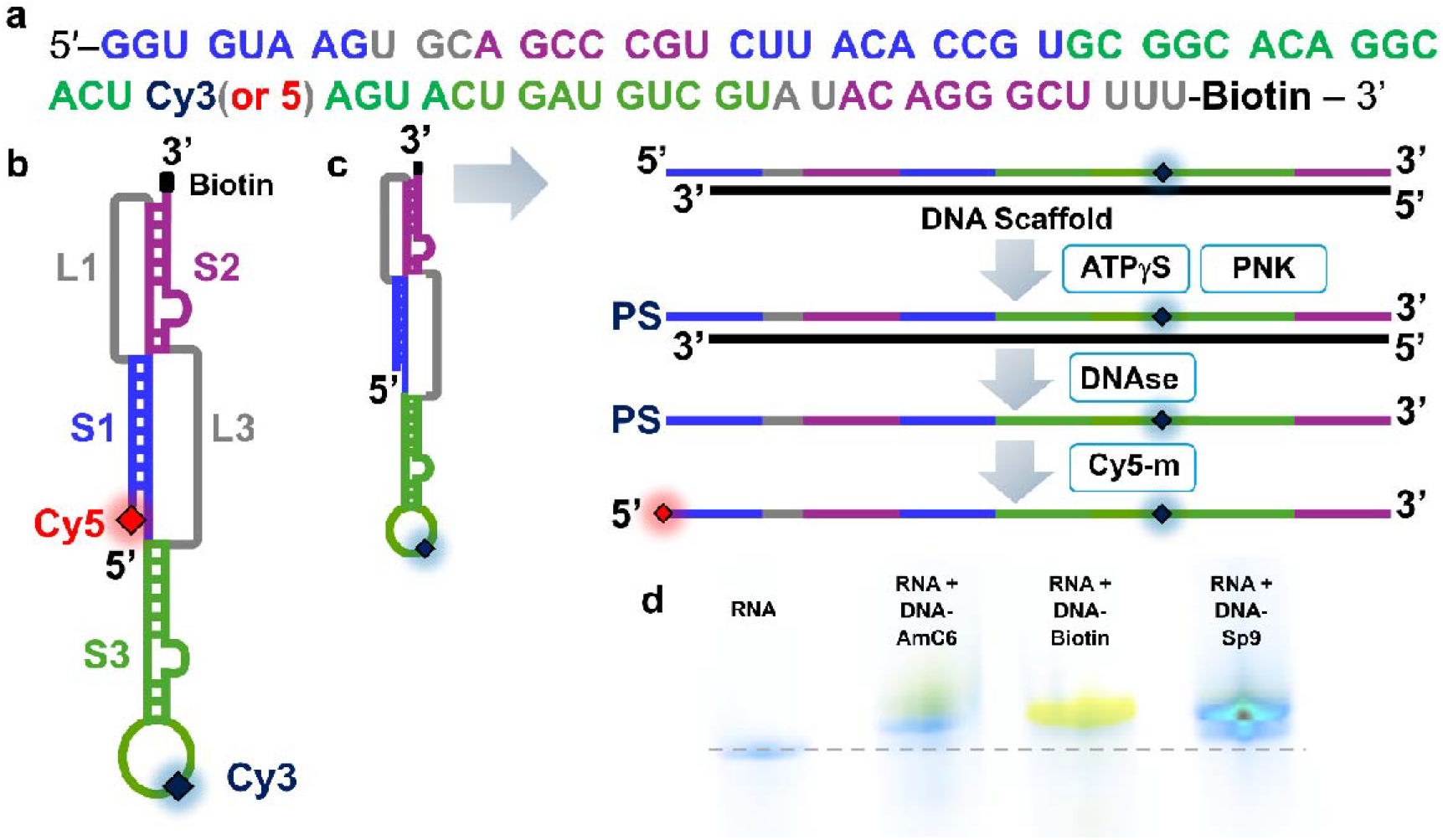
Double labelling of the SARS-CoV-2 RNA pseudoknot via DNA scaffold. (a) Sequence of the pseudoknot with stems and loops color-coded (S1: stem 1, blue; S2: stem 2, purple; S3: stem 3, green; L1/3: loops 1/3, grey). Cy3 and Cy5 labels are shown in navy and red, respectively. (b) Schematic illustration of the RNA secondary structure. (c) Schematic illustration of the labelling process. Initially, a DNA scaffold is added to the internally labelled pseudoknot to form a DNA-RNA duplex, before the addition of ATPγS and polynucleotide kinase (PNK) enable addition of a thiophosphate group (PS) at the 5′ end. After DNA is removed by DNases, Cy5-maleimide (Cy5-m) is added in excess. (d) Fluorescence image of urea PAGE gel before and after the Cy3-labelled pseudoknot was labelled with Cy5-m using DNA scaffolds with AmC6, Biotin and Sp9 5′ end tags. Cy3 emission is shown in blue, and Cy5 emission in yellow. The dashed grey line corresponds to the position of the internally labelled pseudoknot on the gel before labelling.

The following DNA scaffold oligo was also purchased from Integrated DNA technologies and used after RNase-free HPLC purification

5^′^– X AAA AGC CCT GTA TAC GAC ATC AGT ACT AGT GCC TGT GCC GCA CGG TGT AAG ACG GGC TGC ACT TAC ACC – 3^′^.

Here X = 5_′_Biotin, amino C6 (AmC6) or tritheylene glycol (sp9) modification.

### 5_′_ **RNA labelling with DNA scaffold**

6.0μl of internally labelled RNA from a 100μM stock (6 pmol), 2.0 μl 70 mM Tris-HCl, 10 mM MgCl_2_, 5.0 mM Dithiothreitol (DTT) (pH 7.6), 7.0 μl of DNA scaffold from 100μM stock (7 pmol), and 5.0 % spermidine w/v were added to a microcentrifuge tube, heated to 95°C and left to cool for 30 minutes. 1.0 μl of 25 mM ATP_γ_S and 2.0 μl of T4 polynucleotide kinase (3^′^phosphatase minus) (New England Biolabs) were then added and the solution heated to 37°C for a further 1 hour. 2.0 μl RNAse-free DNAse (Thermo), 2.0 μl Duplex DNase (New England Biolabs) and 0.5 μM CaCl_2_ were then added and the solution incubated at 37°C for a further 60 minutes to enable removal of the scaffold. 10 μl of 10 mM Cy3/5 maleimide in dimethyl sulfoxide (DMSO) was subsequently added to 20 μl of this solution and heated to 65°C for 30 minutes. Extraction with pH 4.5 phenol-chloroform was performed and the resulting aqueous phase subjected to ethanol precipitation followed by centrifugation (21,300 x g, 30 minutes, 4°C). Pellets were washed in 70 % ethanol prior to resuspension in ultra-pure water and refolding by heating (80°C, 3 min) and cooling to room temperature. Excess dye was removed by running the solution through a MicroSpin™ G-50 Column (Cytiva). Labelling efficiencies were estimated via absorption spectra generated using a NanoDrop™ 2000/2000c spectrophotometer. Extinction coefficients of 150000 cm^−1^M^−1^ for Cy3 maleimide, 250000 cm^−1^M^−1^ for Cy5 maleimide, 678700 cm^−1^M^−1^ for the Cy3 internal label and 683800 cm^−1^M^−1^ for the Cy5 internal label were used.

### Urea PAGE

The RNA construct under investigation was added to loading buffer (95% (v/v) formamide, 18 mM ethylenediaminetetraacetic acid (EDTA) adjusted to pH 8.0 with KOH, 0.025% sodium dodecyl sulfate, 0.05% bromophenol blue, 0.05% xylene cyanol) and heated to 70°C for 5 minutes. After heating, the solution was placed on ice and loaded onto a 12 % 19:1 acrylimide:bisacrylamide gel containing 7.0 M urea. Electrophoresis was performed in Tris-borate-EDTA buffer (130 mM Tris-HCl pH 7.6, 45 mM boric acid, 2.5 mM EDTA) at constant voltage (300 V) for 30 minutes. Gels were imaged with a Typhoon-5 using 532 (10 mW) and 635 nm (84 mW) excitation lasers, and 570 nm and 670 nm bandpass filters, respectively.

### smFRET Detection

smFRET bursts from freely diffusing RNA molecules in solution were recorded using the smfBox single-molecule fluorescence spectrometer^27^. Briefly, 60 μL droplets of buffer solution (20 mM Tris, pH 7.19) containing 1-5 pM of doubly labelled RNA and NaCl as specified in the main text were placed on a 22 × 22 #1.5 coverslip mounted on a water immersion objective lens (UPLSAPO x60, NA = 1.35). Alternating laser excitation (ALEX) was performed using 515 nm and 635 nm with nominal output powers of 40 mW and 10 mW, respectively and a period of 100 μs. The ALEX cycle involved 45 μs of donor excitation followed by a 5 μs break, 45 μs of acceptor excitation and a final 5 μs time break. Photon arrival times were recorded on two avalanche photodiodes on typical timescales of 60 minutes per sample, at a rate of approximately 1 burst per second.

Analysis was performed using the FRETbursts toolkit^28^. Here, raw photon streams were processed to identify bursts corresponding to donor and acceptor emission from single molecules relative to the background. Apparent FRET efficiencies, E, were estimated using E = I_a_/ (I_a_ + I_d_), where I_d_ and I_a_ are the emission intensities of the donor and acceptor, respectively, obtained from 515 nm excitation. Stoichiometries, S, were calculated according to S = (I_a_ + I_d_)/ (I_a_ + I_d_+ I_a_*), where I_a_* corresponds to the acceptor emission under 635 nm excitation. After background estimation and subtraction, bursts were identified using an all photon sliding window algorithm as previously described^28^. A photon rate of 10 photons (M = 10) and count rate threshold of 16 (F = 16) was applied. A burst size threshold of 20 photons was applied for both donor and acceptor channels. Multimer events were removed from the donor channel by applying an upper limit threshold of 100 photons.

### Accessible Volume Simulations

FRET-restrained positioning and screening software^29, 30^ was used to simulate the accessible volumes of Cy3 and Cy5 on the RNA structure. By combining the PDB files of the static structures and the dimensions of the dyes (length = 17 Å; width = 4.5 Å; dye radius = 3.5 Å), their separation, R, and theoretical FRET efficiency was estimated via 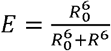, where R is the Förster radius (5.4 nm).

## Results and Discussion

### DNA scaffold-based labelling of the SARS-CoV-2 RNA pseudoknot

We were particularly interested in how the base of stem 1 (S1) affects pseudoknot dynamics. A strong, stable S1 is a hallmark of frameshift-stimulating pseudoknots in coronaviruses^15, 31, 32^ and other viruses^33^. However, the relationship between S1 length and ability to stimulate frameshifting is not well understood. In SARS-CoV-2, stem 3 can make additional interactions with the first C-G base pair of S1, contributing to the formation of “ring-knot” topologies. While this base-pair was observed to form in previous pseudoknot structures^13, 16, 17, 18^ it would nevertheless be the first to break upon an encounter with the ribosome. Given that the full-length wild-type pseudoknot appears to adopt a predominantly static conformation, as observed via cryo-EM^34^, we reasoned that shortening S1 via omission of this base pair might expand the conformational landscape accessible to the RNA. We therefore omitted the 5^′^ cytosine from our construct to probe how disruption at the S1 base influences folding dynamics.

We used maleimide click chemistry to conjugate the FRET acceptor Cyanine 5 (Cy5)^26^ onto molecules harbouring an internal FRET donor (Cyanine 3, Cy3) (**Figure 1a**). By placing the internal label at the apex of stem 3 and another on the 5^′^ end (**Figure 1b**), we hypothesized that the locations would enable smFRET to sensitively report on long-range conformational changes associated with mobility of pseudoknot tertiary elements. Initially, we explored a conventional maleimide labelling strategy, using T4 polynucleotide kinase (PNK) to transfer a thiophosphate group from ATPγS onto the 5^′^ end of the molecule. However, direct incubation of the PNK and ATPγS treated molecule with a molar excess of Cy3 was ineffective, providing a sub-optimal labelling efficiency of ∼ 3 % as estimated by absorption spectroscopy. Based on the available crystal and cryo-EM structures, we hypothesized that the low labelling efficiency was likely due to inaccessibility of the 5^′^ end during the enzymatic labelling step, preventing PNK activity^35^.

To circumvent this challenge, we took inspiration from splint ligation^36, 37^. A DNA scaffold was designed to anneal to the RNA molecule, effectively removing all secondary structure, and creating a three nucleotide 5^′^ overhang of single-stranded RNA to more effectively enter the PNK active site^35^. To prevent unwanted PNK activity on the DNA scaffold, we synthesised the DNA with 5^′^ modifications to create steric hindrance with the enzyme active site. To this end, we evaluated the effects of DNA variants containing 5^′^ biotin, Amino (AmC6) and triethylene glocol (Sp9) modifications. After PNK and ATPγS incubation, the combined use of Duplex DNAse and DNAse I was introduced to (i) endonucleolytically cleave the DNA when hybridised to RNA and (ii) remove the fragments. Cy5-maleimide was subsequently conjugated to a thiophosphate group introduced by PNK at the 5^′^ end of the RNA. The protocol to achieve double labelling is schematically shown in **Figure 1c**, and detailed instructions are provided in the **Methods**. Urea PAGE confirmed the presence of doubly labelled species (**Figure 1d**), and absorption spectroscopy indicated over a 7-fold improvement in labelling efficiency (∼23 %). The three 5^′^ DNA modifications – biotin, AmC6 and Sp9 – were directly compared on the gel (**Figure 1d**) to assess their impact on labelling efficacy. We note that replacing the 5^′^ biotin modification with an AmC6 or Sp9 spacer led to reduced labelling efficiencies of ∼6 % and 16 %, respectively, likely due to differences in how these 5^′^ modifications influence duplex stability or geometry.

### smFRET characterisation of the SARS-CoV-2 RNA pseudoknot in solution

Having confirmed the presence of both donor and acceptor labels on the pseudoknot structure, smFRET measurements were acquired on freely diffusing molecules in solution using confocal microscopy. Here, diffusion of doubly labelled molecules was monitored via fluorescence emission as they transited through a confocal excitation volume. In this case we used alternating laser excitation (ALEX) to enable excitation of the Cy3 and Cy5 fluorophores separately, enabling millisecond timescale photon bursts of fluorescence in both donor and acceptor detection channels (**Figure 2a**). Photon counts from both channels (**Figure 2b**) were then converted into estimates of the apparent FRET efficiency, E, and stoichiometry, S, of labelling for each molecule. In line with our labelling efficiency estimates, we detected a high proportion of donor-only labelled molecules compared to doubly labelled species (**Figure 2c**). However, the ALEX approach allowed us to separate both donor-only and acceptor-only species from the doubly labelled population^27^, opening a platform for analysis of the FRET-active species (**Figure 2d**) as a function of local environment.

**Figure 2.**
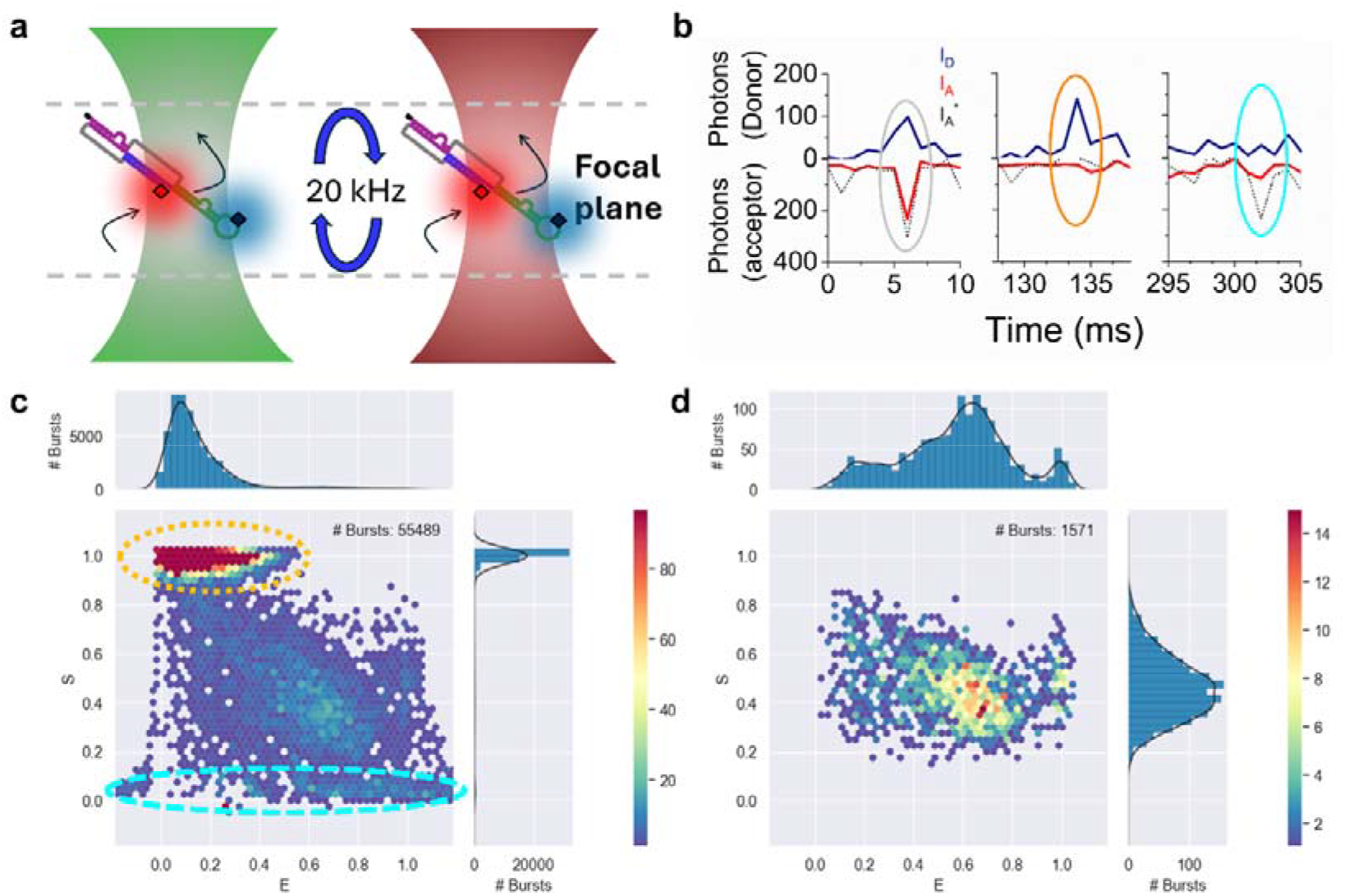
smFRET detection of freely diffusing Cy3-Cy5 labelled RNA pseudoknots. (a) Schematic illustration of the setup. Single RNA molecules freely diffuse through a confocal excitation volume on millisecond timescales. 515 nm and 635 nm are alternated (20 kHz) to ensure multiple excitation cycles of Cy3 and Cy5 for each diffusing molecule. Arrows indicate diffusion of the molecule through the confocal volume. (b) Representative time traces from a smFRET experiment. Fluorescently labelled RNA molecules freely diffuse through the confocal volume emitting bursts of fluorescence. Here, individual photons are collected from the donor (I_D_, navy) and acceptor (I_A_, red) under 515 nm excitation, and from the acceptor under 635 nm excitation (I_A_*, black), enabling discrimination of doubly labelled species (left panel), those containing only a donor molecule (middle) and those only containing only an acceptor (right panel). The grey, orange and cyan ovals highlight the regions of interest. (c) 2D apparent FRET efficiency and stoichiometry plot. RNA molecules containing only Cy3 appear with low E and high S (orange dashed region), whereas Cy5-labelled molecules (no Cy3) appear with low S (cyan dashed region). (d) Doubly labelled molecules (N = 1571), appearing with intermediate S, extracted from the data shown in (c). Solution conditions: 20 mM Tris, pH 7.6.

### The SARS-CoV-2 frameshift stimulatory element displays salt-dependent conformational heterogeneity

In the absence of NaCl, we identified a broad distribution of FRET efficiency values spanning the full available range (0 – 1) (**Figure 3a**), pointing towards flexibility within the molecule. A multiple-Gaussian model applied to the experimental data revealed four populations with FRET states centred on 0.21 (FWHM = 0.18) (state 1), 0.42 (FWHM = 0.11) (state 2), 0.63 (FWHM = 0.22) (state 3) and and 0.99 (FWHM = 0.04) (state 4). The relative proportion in each state, evaluated by the area of each Gaussian distribution was 18 %, 12 %, 64 % and 6 %, respectively. In the presence of 150 mM NaCl, however, states 1 and 4 diminished, and the distribution shifted towards states 3 (FWHM = 0.23) and 2 (FWHM = 0.28), with the majority (77 %) residing in state 3 (**Figure 3a**). Indeed, analysis of the relative difference between the two distributions indicated substantial differences in population across all 4 states (**Figure 3b**), further pointing to NaCl-dependent conformational shifts. Similar observations were made when the RNA molecule was internally labelled with Cy5, and 5^′^ labelled with Cy3, giving confidence that the observed variations were due to NaCl-induced conformational rearrangements (**Figures 3c, d**).

**Figure 3.**
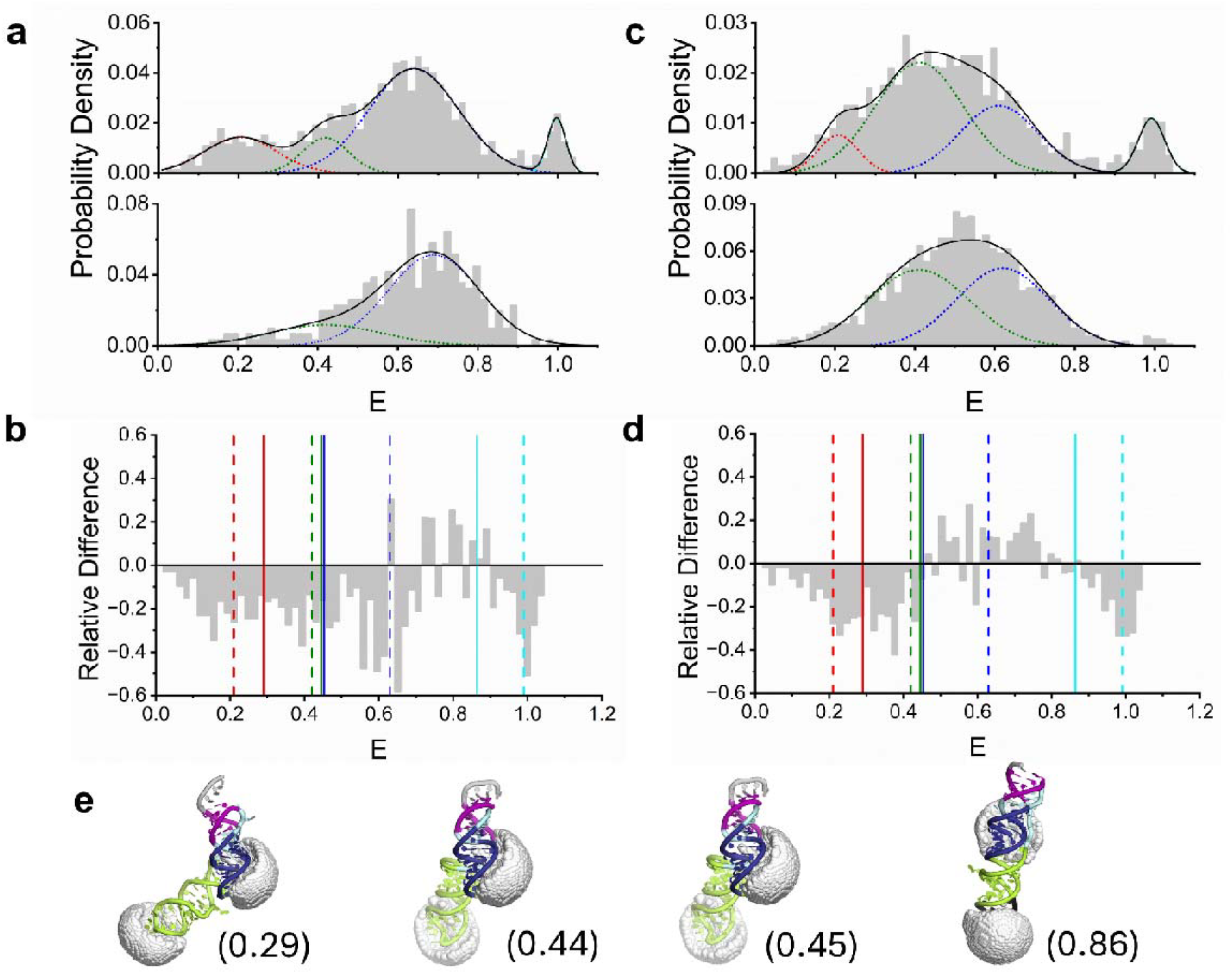
NaCl induces the structural remodelling of freely diffusing RNA pseudoknots in solution. (a) smFRET population distributions obtained from freely diffusing RNA pseudoknots internally labelled with Cy3, and 5′ labelled with Cy5 in the absence (top panel, N = 1571) and presence (bottom panel, N = 312) of 150 mM NaCl. A multi-Gaussian fit (black, χ^2^ = 0.93) comprising states 1 (E = 0.21, red), 2 (0.42, green), 3 (0.63, blue) and 4 (0.99) was applied in the absence of NaCl, and a bi-Gaussian fit (χ^2^ = 0.90) comprising states 2 and 3 were applied in the case of 150 mM NaCl. (b) Relative difference between the 150 mM and 0 mM NaCl distributions. Centres of the Gaussians are indicated as dashed lines. Solid lines correspond to theoretical FRET efficiency predictions from AV simulations. (c) Corresponding smFRET distributions obtained from pseudoknots internally labelled with Cy5 and 5′ labelled with Cy3 in the absence (top panel, N = 1266) and presence (bottom panel, N = 2009) of 150 mM NaCl. A multi-Gaussian fit (black, χ^2^ = 0.92) was applied in the absence of NaCl, and a bi-Gaussian fit (χ^2^ = 0.95) was applied in the case of 150 mM, each comprising states as described in (a). (d) The relative difference between distributions is also shown with Gaussian centres indicated as dashed lines. Solid lines correspond to theoretical FRET efficiency predictions from AV simulations. (e) AV simulations of the molecule from static predictions (PDB IDs from left to right: 707Y, 6XRZ, 7MKY and 6XRZ) containing Cy3 (internal) and Cy5 (5′ end). Stem 1, stem 2 and stem 3 are shown in blue, purple and green, respectively, and accessible volumes are shown in grey. Numbers in brackets correspond to the theoretical FRET efficiency predictions.

It is interesting to note that differences in the FRET distributions between the two dye-labelling configurations were observed. For example, in the case where the molecule contained a Cy5 internal label, state 2 was more pronounced by comparison (**Figure 3c**). While further work is required to fully elucidate this observation, it is worth noting that differences between the FRET distributions could arise from the orientation sensitivity of FRET as well as possible dye-RNA interactions. Although the FRET efficiency primarily depends on inter-dye distance, it is also influenced by the relative orientation of the donor and acceptor dipoles, which can differ depending on dye position and local structural changes. The internal dye, for instance, may be more constrained than the dye at the 5^′^ end, as a consequence of being attached at two points, and the labels may exhibit subtly different rotational freedom depending on their position and size, thereby altering the absolute FRET efficiencies recovered. Variations in electrostatic interactions may also shift the equilibrium, explaining why state 2 appears more prominent in one labelling scheme versus the other. Nevertheless, under both conditions we observed (i) a clear distribution of states, pointing towards conformational heterogeneity of the pseudoknot in solution, and (ii) removal of low- and high-FRET populations, and narrowing of the distributions upon NaCl addition, suggesting that the local salt content plays an important role in modulating the conformational landscape.

When a direct comparison was made between the experimentally determined FRET efficiencies and accessible volume simulations (**Figure 3e**) which recover the absolute distances between donor and acceptor pairs conjugated to the known structures, we hypothesise that intermediate FRET states ranging from 0.2-0.6 are consistent with combinations of previously determined L-shaped^13, 38^and fully twisted linear^17^ conformations where theoretical dye separation distances of 61 Å, 52 Å and 53 Å were predicted. We note that the predictions were identical under both labelling schemes as they are based on linker length. By contrast, the high FRET state (state 4) observed in the absence of NaCl and under both scenarios is consistent with a shortened version of the untwisted linear conformation which predicted a FRET efficiency of 0.86 and dye separation distance of 34 Å^18^ (**Figure 3b, d**). In other words, we speculate that the 5^′^ end of the molecule regulates the conformational landscape, and that addition of NaCl leads to stabilisation of the base-pairs in stem 3 and reduction of untwisted structure. While further experiments are necessary to fully validate this hypothesis, the broad distributions observed under each condition tested indicates that the pseudoknot with a single C-G deletion in S1, likely explores a NaCl-dependent heterogeneous conformational landscape in solution, that could also involve dynamic transitions between states. This work, therefore, serves as a starting point for exploring the environmental variables and factors which influence the pseudoknot structure in solution, and could be extended towards evaluation of the structure in the presence of a wide variety of local environmental factors and ligands.

## Conclusions

We have reported a new double-labelling strategy and the smFRET characterisation of freely diffusing SARS-CoV-2 RNA pseudoknot frameshifting elements in solution. By using a 5^′^-modified DNA scaffold to remove RNA secondary structure, we were able to increase PNK activity enough to facilitate dye conjugation at an acceptable efficiency, enabling quantification of conformational heterogeneity via smFRET population distributions. We identify that the molecule, disrupted at the first C-G base pair in S1, samples a multiplicity of NaCl-dependent states in solution. This represents a proof-of-concept for our reporter design, which will be used to evaluate the relative importance of base pairing at every S1 position in future work. This study therefore provides a starting point for addressing a range of questions, including elucidating the conformational landscape of the molecule as a function of environment, assessing interactions induced by binding partners, and evaluating the kinetic switching between conformational states. In short, this work forms the basis of a new method for interrogating the structure, dynamics and heterogeneity of complex RNA biomolecules at the single-molecule level, which in turn could provide information that that is otherwise hidden from structural methods such as NMR, X-ray crystallography and cryo-EM. We suggest this labelling strategy will be a useful addition to the RNA toolbox, alongside established techniques such as co-transcriptional labelling, 3^′^ end oxidation and splint ligation.

## Acknowledgements

We thank Dr Mahmoud Abdelhamid (University of Sheffield, UK) for help with initial exploratory work. S. D. Q., M. C. L. and C. H. H. thank the Engineering and Physical Sciences Research Council (EPSRC) Impact Accelerator (EP/X525856/1) for support. C.H.H. is supported by a Sir Henry Dale Fellowship (221818/Z/20/Z) from the Wellcome Trust and the Royal Society. C.H.H. is also supported by the Lister Institute of Preventive Medicine. S. P. G. is supported by an EPSRC studentship (EP/T518025/1). M. L. is supported by EPSRC (EP/W024063/1 and EP/Y000501/1) and the Biotechnology and Biological Sciences Research Council (BB/W000555/1). J.K.B. is supported by a Medical Research Council DiMeN PhD studentship (MR/W006944/1). The authors also acknowledge Dr Andrew Leech, Dr Katy Cornish, Dr Peter O’Toole and Dr Alex Payne-Dwyer for technical support within the York Bioscience Technology Facility.

